# Synonymous codon substitutions perturb co-translational protein folding *in vivo* and impair cell fitness

**DOI:** 10.1101/666552

**Authors:** Ian M. Walsh, Micayla A. Bowman, Iker F. Soto, Anabel Rodriguez, Patricia L. Clark

## Abstract

In the cell, proteins are synthesized from N- to C-terminus and begin to fold during translation. Co-translational folding mechanisms are therefore linked to elongation rate, which varies as a function of synonymous codon usage. However, synonymous codon substitutions can affect many distinct cellular processes, which has complicated attempts to deconvolve the extent to which synonymous codon usage can promote or frustrate proper protein folding *in vivo*. Although previous studies have shown that some synonymous changes can lead to different final structures, other substitutions will likely be more subtle, perturbing predominantly the protein folding pathway without radically altering the final structure. Here we show that synonymous codon substitutions encoding a single essential enzyme lead to dramatically slower cell growth. These mutations do not prevent active enzyme formation; instead, they predominantly alter the protein folding mechanism, leading to enhanced degradation *in vivo*. These results support a model where synonymous codon substitutions can impair cell fitness by significantly perturbing co-translational protein folding mechanisms, despite the chaperoning provided by the cellular protein homeostasis network.

**Significance:** Many proteins that are incapable of refolding *in vitro* nevertheless fold efficiently to their native state in the cell. This suggests that more information than the amino acid sequence is required to properly fold these proteins. Here we show that synonymous mRNA mutations can alter a protein folding mechanism *in vivo*, leading to changes in cellular fitness. This work demonstrates that synonymous codon selection can play an important role in supporting efficient protein production *in vivo*.

## Introduction

Synonymous codon substitutions alter the mRNA coding sequence but preserve the encoded amino acid sequence. For this reason, these substitutions were historically considered to be phenotypically silent and often disregarded in studies of human genetic variation (1, 2). In recent years, however, it has become clear that synonymous substitutions can significantly alter protein function *in vivo* through a wide variety of mechanisms that can change protein level (3–5), translational accuracy (6, 7), secretion efficiency (8, 9), the final folded structure (1, 10–12) and post-translational modifications (13). The full range of synonymous codon effects on protein production is, however, still emerging and much remains to be learned regarding the precise mechanisms that regulate these effects.

One effect of synonymous codon substitutions long proposed but with scant evidence to support its significance *in vivo* is perturbations to co-translational folding mechanisms. In general, rare synonymous codons tend to be translated more slowly than their common counterparts (14–17). Moreover, rare synonymous codons tend to appear in clusters, creating broader patterns of codon usage (18), many of which are conserved through evolution (19–21). The folding rates of many protein secondary and tertiary structural elements are similar to their rate of synthesis (22, 23), lending conceptual support to the hypothesis that even subtle changes in elongation rate could alter folding mechanisms (24). In theory, reducing the rate of translation elongation by synonymous common-to-rare codon substitutions could provide the N-terminal portion of a nascent protein with more time to adopt a stable tertiary structure before C-terminal portions are synthesized and emerge from the ribosome exit tunnel (25–27). Depending on the specific native structure of the encoded protein, such extra time could be either advantageous or detrimental to efficient folding (28). However, cells contain an extensive network of molecular chaperones to facilitate the folding of challenging protein structures, including several that associate with nascent polypeptide chains during translation (29–33). Thus, it remains unclear whether a synonymous codon-derived alteration to elongation rate and co-translational folding mechanism could be sufficiently perturbative to rise above the buffering provided by the cellular chaperone network.

Here we show that synonymous codon changes in the coding sequence of an enzyme essential for *E. coli* growth can have a dramatic impact on cell growth. We tested a variety of mechanistic origins for this growth defect, including changes to the folded protein structure, expression level, enzymatic activity, mRNA abundance and/or activation of a cell stress response. Our results are consistent with synonymous substitutions altering the pattern of translation elongation, which alters the co-translational folding mechanism and leads to the formation of a folded, active structure that is more susceptible to degradation. These results demonstrate that changes to synonymous codon usage can significantly affect protein folding *in vivo*, rising above the chaperoning capacity provided by the cellular protein homeostasis network. Synonymous codon usage may therefore have broad implications for effective protein design and the interpretation of disease-associated synonymous mutations.

## Results

### *Synonymous codon substitutions impair* E. coli *growth rate*

To develop a system to test connections between synonymous codon usage, co-translational folding and cell fitness, we used chloramphenicol acetyltransferase (CAT), a water-soluble, homotrimeric *E. coli* enzyme with a complex tertiary structure (34) (**Fig. 1**). A landmark early study showed that synonymous codon substitutions near the middle of the coding sequence (**Fig. 2a, S1**) led to lower specific activity for CAT synthesized by *in vitro* translation (11). CAT is essential for *E. coli* growth in the presence of chloramphenicol (cam) (35), which enabled us to use growth rate with cam as a convenient fitness assay. Furthermore, because CAT is not part of an operon or regulatory network, we hypothesized that it would be unlikely for feedback regulation of other genes to mask the effects of CAT synonymous codon changes on enzyme function (36). Crucially, although CAT cannot be refolded to its native structure after dilution from chemical denaturants, the native structure is resistant to unfolding up to 80°C (**Fig. S2**), suggesting that folding intermediates populated during and after protein synthesis are crucial for efficient folding, as once the native structure has been attained it is not likely to populate the unfolded state over a typical bacterial lifespan.

**Figure 1.**
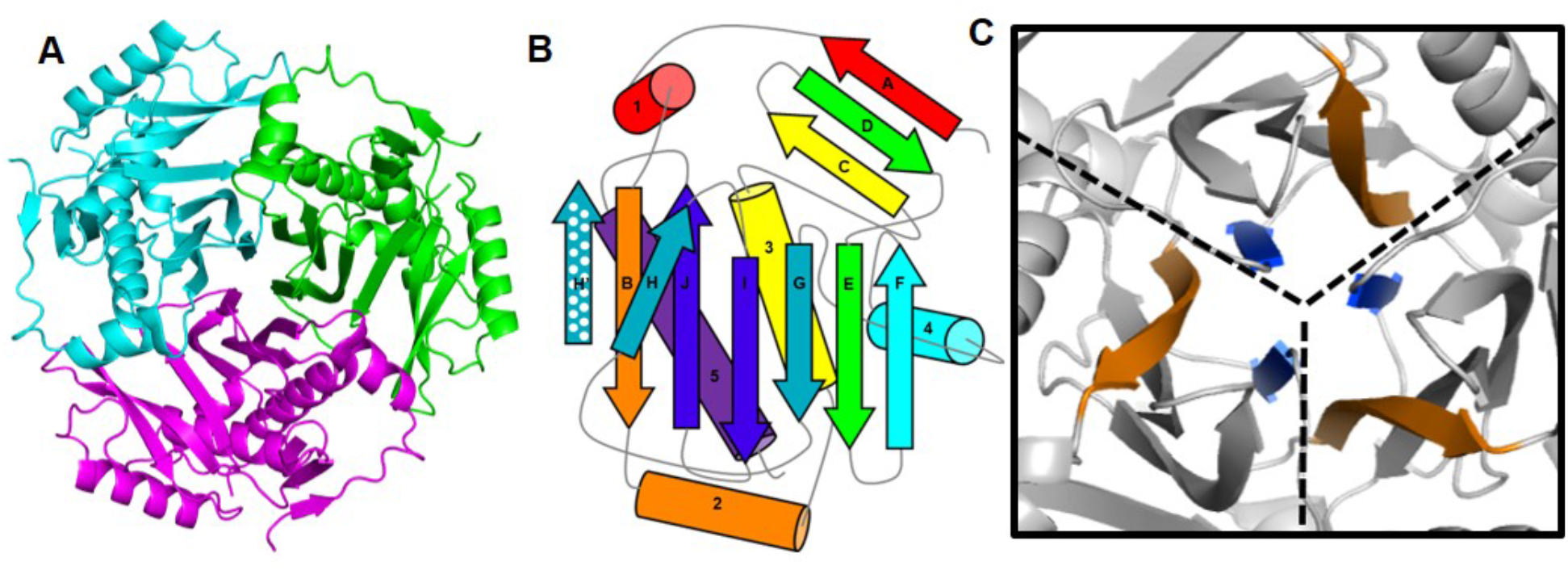
Chloramphenicol acetyltransferase (CAT) has a complex tertiary and quarternary structure. (**A**) Ribbon diagram depicting the native homotrimeric structure (PDBID: 3CLA) (34). (**B**) Schematic representation of the complex topology of the CAT monomer structure. Secondary structure elements are shown in rainbow order. Polka dots indicate the H β-strand in the central β-sheet contributed from an adjacent monomer. (**C**) Close up of the trimer interface, with the B and H β-strands in the central β-sheets colored as in panel (B). Dashed lines indicate approximate monomer boundaries.

We transformed *E. coli* with a plasmid encoding the previously described synonymous CAT coding sequence variant (11) under a titratable promoter but detected no discernable difference in growth versus *E. coli* producing CAT from the wild type (WT) coding sequence (**Fig. 2b, S3a**). However, compared to WT-CAT, this synonymous construct contains a larger number of common codons (**Fig. 2a**), which leads to increased protein accumulation due to an overall faster translation elongation rate (11, 16, 25). Consistent with this, we detected more CAT in *E. coli* transformed with this coding sequence enriched in common codons (**Fig. 2c**). We hypothesized that this higher intracellular CAT concentration could mask a defect in specific activity. To test this, we used a Monte Carlo simulation method (18, 37) (see Supplemental Methods) to design and select an alternative synonymous CAT coding sequence, Shuf1. In Shuf1, the local synonymous codon usage patterns are very different from the WT coding sequence but the global codon usage frequencies are very similar (**Fig. 2a, S1**), which we predicted would lead to the synthesis of a WT-like amount of CAT. To avoid known effects of 5’ synonymous codon substitutions on translation initiation (5, 38–41), the first 46 codons of Shuf1 are identical to the WT coding sequence. Consistent with our prediction, *E. coli* produced CAT from the Shuf1 coding sequence at levels indistinguishable from WT-CAT (**Fig. 2c**). However, cells expressing Shuf1-CAT grew more slowly than cells expressing WT-CAT (**Fig. S3a**).

**Figure 2.**
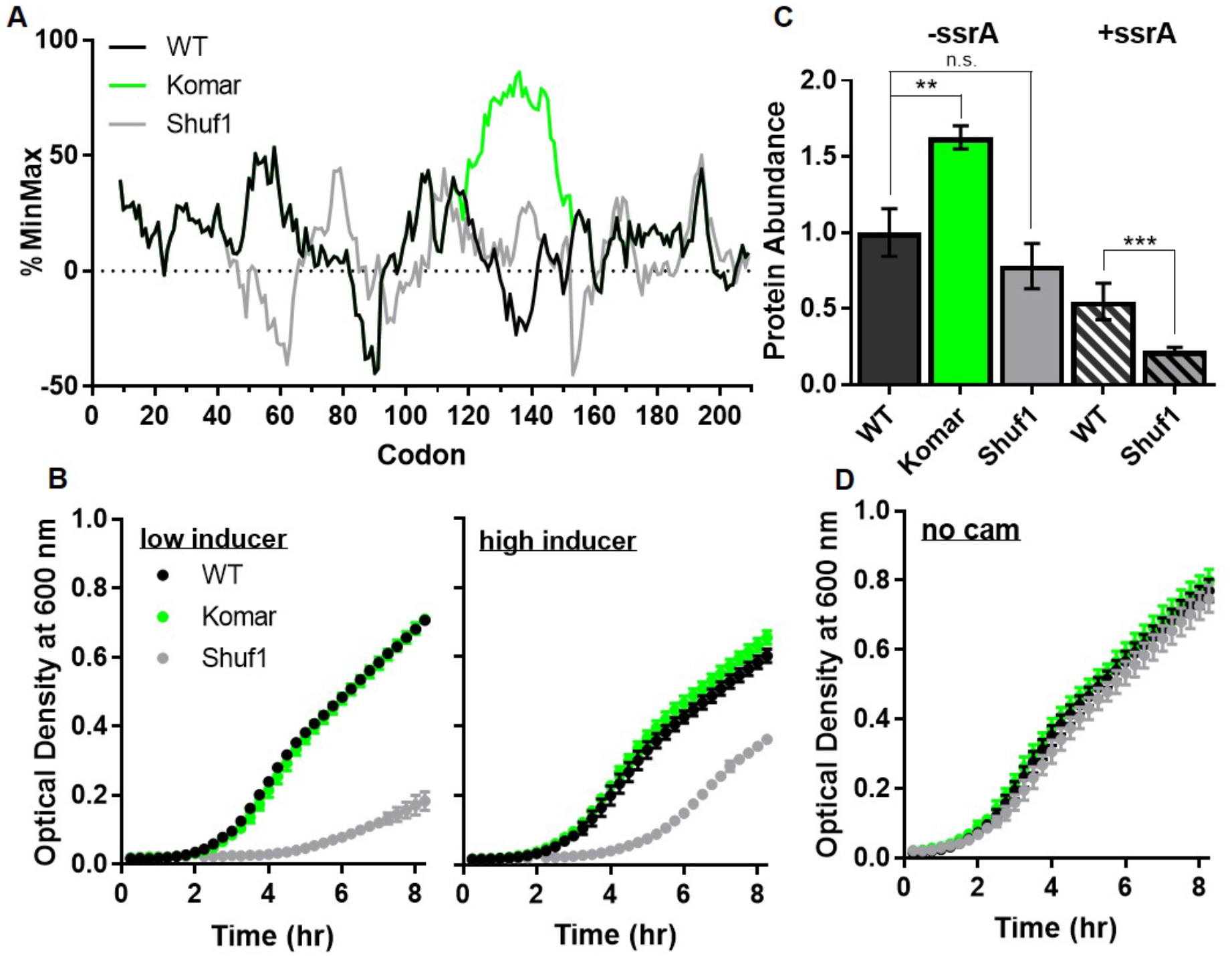
CAT encoded by the synonymous Shuf1 sequence leads to impaired *E. coli* growth in the presence of chloramphenicol (cam). (**A**) Relative codon usage in WT (black), Komar (11) (green), and Shuf1 (gray) CAT coding sequences. Positive values correspond to clusters of common codons and negative values represent clusters of rare codons, calculated over a sliding window of 17 codons (37). (**B**) Growth curves of *E. coli* expressing ssrA-tagged CAT variants challenged with cam under low (200 ng/mL) or high (1,600 ng/mL) concentrations of inducer. (**C**) Relative abundance of untagged (solid bars) or ssrA-tagged (hatched bars) CAT accumulated in cells determined by quantitative western blotting of cell lysates. (**D**) Growth curves in the absence of cam. In all figures, data points represent the mean ± SD of at least three independent experiments; ** p < 0.01, *** p < 0.001, Welch’s t-test.

We hypothesized that we could further exacerbate the observed Shuf1-CAT growth defect by adapting a strategy developed by Hilvert and coworkers to couple subtle changes in enzyme function to *E. coli* growth rate (42). This strategy involves encoding a ClpXP recognition tag (ssrA) at the C-terminus of the protein of interest, selectively enhancing its degradation by the *E. coli* AAA+ protease ClpXP and leading to correspondingly lower intracellular protein concentrations. Addition of the ssrA tag did not affect CAT structure, stability or specific activity (**Fig. S3b-d**) but did lead to a dramatic growth defect for *E. coli* expressing Shuf1-CAT, versus ssrA-tagged WT-CAT, in the presence of cam (**Fig. 2b**). This defect also led to a lower minimum inhibitory concentration for *E. coli* expressing Shuf1-versus WT-CAT (**Fig. S3e**).

### Neither the Shuf1-CAT mRNA nor protein is inherently toxic

A major challenge of all *in vivo* experiments is discerning the precise origin of an observed effect. For example, a recent study indicated that synonymous codon substitutions can lead to toxicity at the mRNA level even in the absence of protein production (43). To test whether production of the Shuf1-CAT ssrA mRNA and/or protein is inherently toxic, we compared the growth of *E. coli* expressing WT or Shuf1-CATssrA in the absence of cam. These growth curves were indistinguishable (**Fig. 2d**), indicating that the Shuf1 defect is specifically related to impaired CAT enzyme function. Moreover, in the presence of cam the growth defect was partially suppressed at higher inducer concentrations (**Fig. 2b**), contrary to the larger growth defect expected if the Shuf1-CATssrA mRNA and/or protein were inherently toxic.

To test whether Shuf1-CAT expression induces a general cell stress response, we used mass spectrometry to compare the abundances of 1277 proteins in *E. coli* expressing ssrA-tagged CAT from either the WT or Shuf1 coding sequence. There was no significant difference detected in the level of most proteins, including known stress-associated molecular chaperones and proteases (**Fig. 3**). Taken together, these results support a model where the Shuf1-CAT growth defect is due to a direct defect in active CAT protein production, rather than an indirect effect on other cell functions.

**Figure 3.**
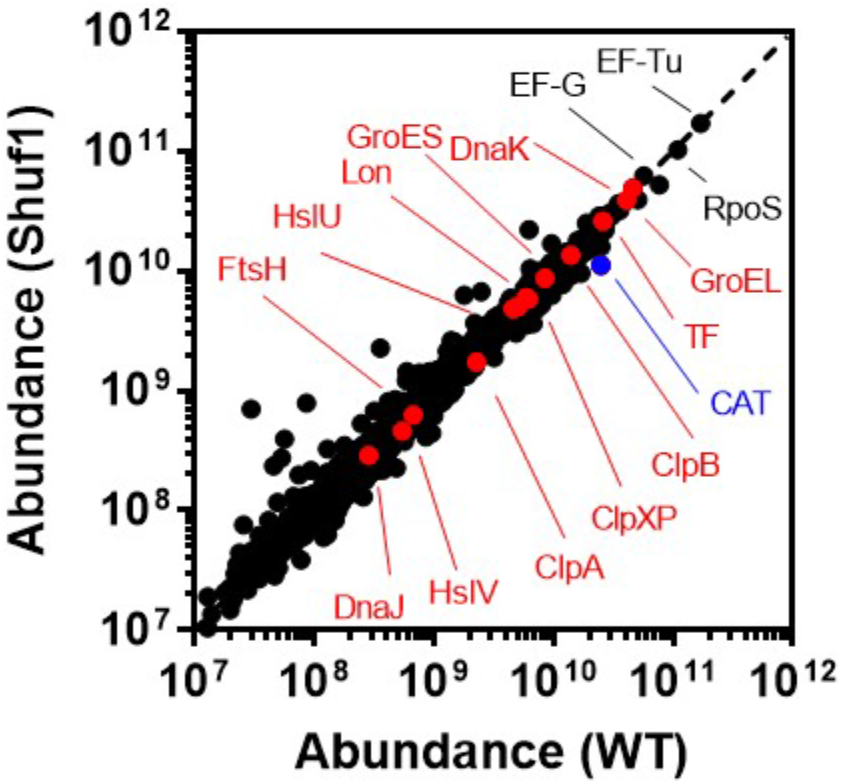
Translation of CAT using Shuf1 coding sequence does not significantly perturb the *E. coli* proteome. Relative abundance of *E. coli* proteins upon expression of WT or Shuf1 CAT. Twelve *E. coli* molecular chaperones and AAA+ ATPases are shown in red; 1264 other *E. coli* proteins are shown in black. No significant upregulation of chaperones or ATPases was observed for *E. coli* expressing Shuf1.

### Shuf1 coding sequence does not adversely affect mRNA concentration

We noticed that addition of the ssrA tag led to a larger reduction in intracellular accumulation for CAT produced from the Shuf1 versus WT coding sequence (**Fig. 2c**, hatched bars). To determine whether this decrease in Shuf1-CAT was due to a defect arising from Shuf1 transcription and/or mRNA half-life, versus a translation-related defect, we quantified the levels of WT and Shuf1 mRNA. These levels were indistinguishable (**Fig. S4a**). Together with the indistinguishable levels of WT- and Shuf1-CAT protein accumulation in the absence of the ssrA tag (**Fig. 2c**, filled bars), these results suggest a model where the Shuf1 synonymous codon changes affect intracellular CAT concentration at the translational level, likely due to a greater susceptibility of the Shuf1-CAT protein to degradation (**Fig. 4**).

**Figure 4.**
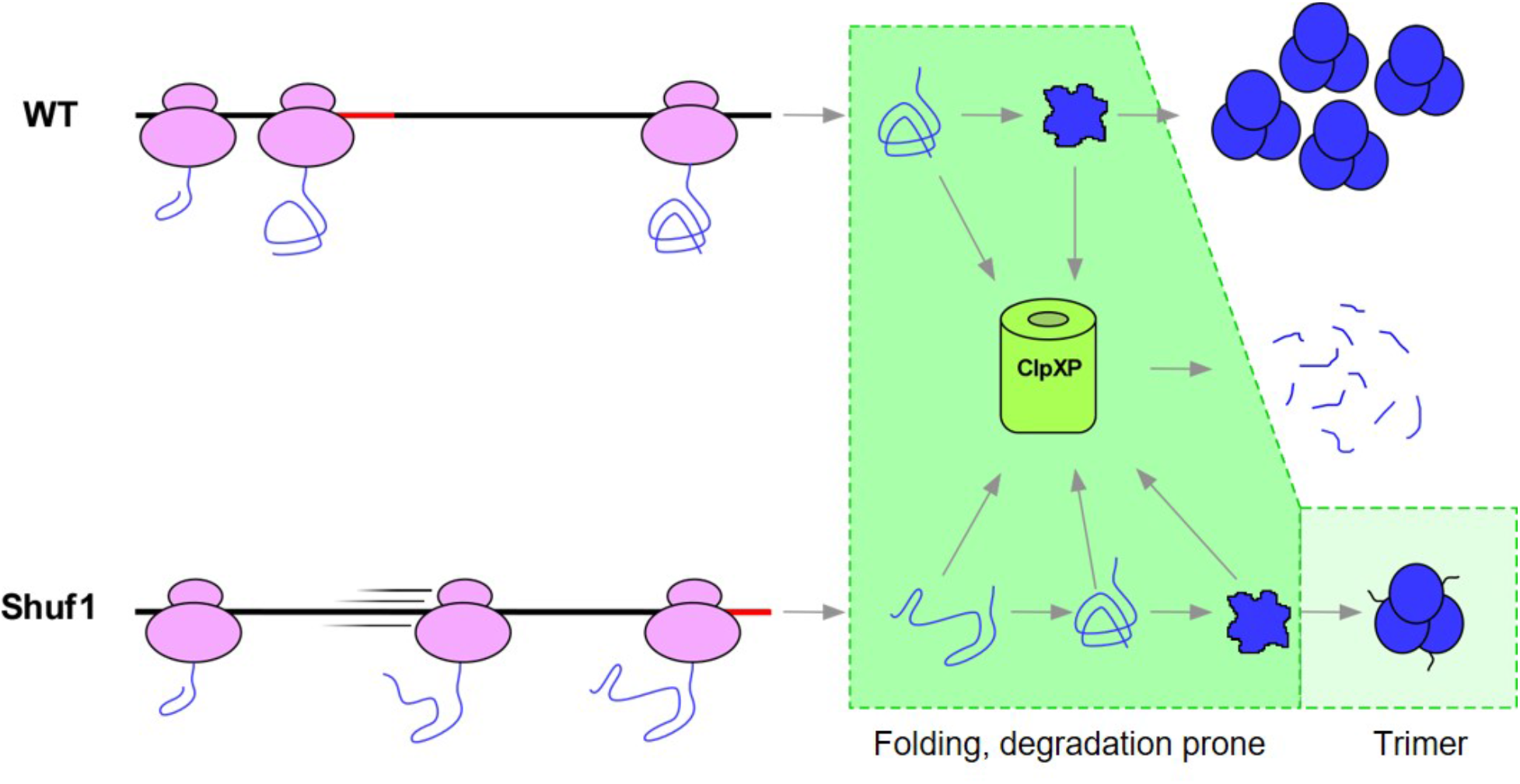
Proposed model for the effects of synonymous CAT codon substitutions on ssrA-tagged CAT folding and cell fitness. Synonymous changes in the Shuf1 coding sequence alter the local rate of translation, affecting the conformation of CAT co-translationally and persisting after release of the nascent protein from the ribosome. These altered Shuf1 folding intermediates are more susceptible to degradation by ClpXP than intermediates populated during and after translation of the WT coding sequence. Some Shuf1-CATssrA proteins evade degradation and eventually fold to an active conformation that is also more susceptible to degradation than WT-CATssrA.

### ClpX deletion indicates ClpXP is major source of Shuf1-CATssrA growth defect

If the Shuf1 codon-dependent growth defect is due to more efficient degradation of ssrA-tagged Shuf1-CAT by cellular proteases, specifically ClpXP, deleting ClpX would be expected to ameliorate the growth defect *in vivo*. ClpXP is the major *E. coli* protease responsible for degrading ssrA-tagged polypeptides under log-phase growth (44, 45). In general, less stably-folded proteins are more susceptible to degradation by ClpXP than more stable substrates (46–48), presumably because less energy is required for ClpX to unfold unstable protein structures and expose the polypeptide chains to the ClpP protease active sites (49). To test whether ClpXP degradation is the key mechanism impairing growth when *E. coli* expresses CAT from the Shuf1 coding sequence, we induced expression of WT and Shuf1 CAT in an *E. coli* W3110 derivative that lacks ClpX (46, 50) and compared growth in this background to the parent strain W3110, in the presence of cam. ClpX deletion enhanced growth only of cells expressing ssrA-tagged CAT from the Shuf1 coding sequence (**Fig. 5a**). Likewise, omission of the ssrA tag enhanced growth only for *E. coli* expressing ClpX; there was no effect on cells lacking ClpX (**Fig. 5b**). These results confirm that the major effect of the Shuf1 synonymous codon substitutions is enhanced degradation of ssrA-tagged CAT by ClpXP.

**Figure 5.**
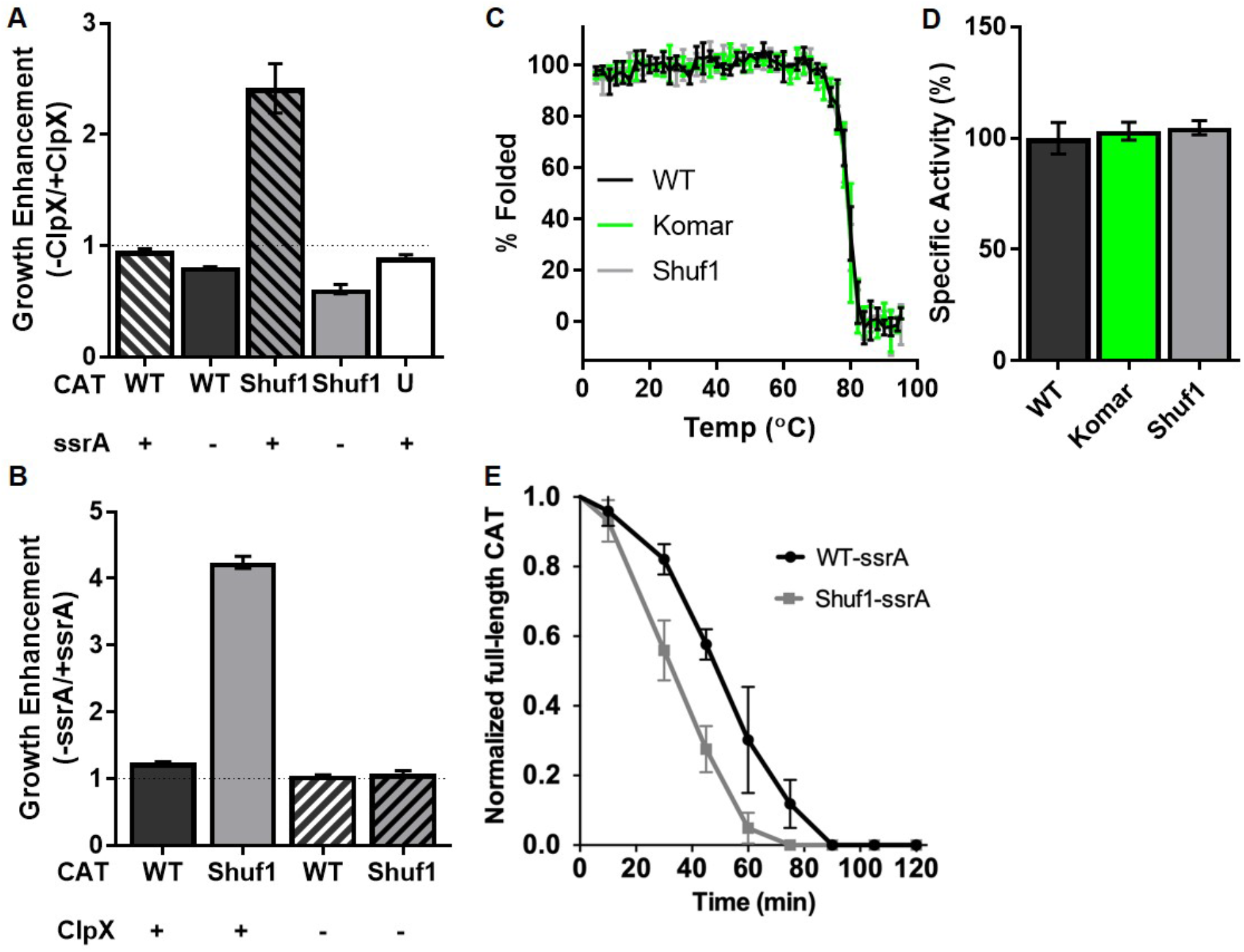
Shuf1 CAT is more susceptible to ClpXP degradation than WT CAT, despite several other indistinguishable characteristics. (**A-B**) Selective effects of ssrA-tagging and ClpX deletion on the Shuf1 growth defect. (**A**) In the ClpX deletion strain (W3110 *ΔclpX*), a large increase in growth rate relative to the parent strain is observed only for ssrA-tagged Shuf1. Other constructs grow slightly slower in the absence of ClpX. U, uninduced cell culture. (**B**) Cell growth data from panel (A) plotted to highlight the impact on growth rate of removing the ssrA tag. Omitting the ssrA tag has no effect on growth in the ClpX knockout (hatched bars). In the presence of ClpX (filled bars), there is a much larger increase in growth upon ssrA tag deletion for Shuf1 than WT, indicating Shuf1 is more susceptible to ClpXP degradation than WT. (**C**) Thermal denaturation of CAT monitored by far-UV CD spectroscopy at 205 nm. (**D**) Acetyltransferase activity of purified, native CAT, normalized to WT. (**E**) *In vitro* ClpXP degradation of native, purified, ssrA-tagged CAT trimers. CAT-ssrA bands are indicated along with the components of the assay (ClpXΔN_3_: ClpX trimer without N terminal domains [43] CPK: creatine phosphokinase, and ClpP). In all panels, data points represent the mean ± SD; n=3 biological replicates.

### Native WT- and Shuf1-CAT proteins are subtly different

Synonymous codon substitutions can lead to a wide range of effects on the encoded protein, including changes to translational fidelity (decoding accuracy) (6) and the native structure (1, 10, 12, 17). As a next test of the mechanism by which Shuf1 codon changes alter cell growth rate, we compared the CAT proteins produced from the WT and Shuf1 coding sequences. In both cases, CAT was detected only in the soluble fraction of the cell lysate (**Fig. S4b**), indicating the Shuf1 growth defect is not due to CAT aggregation. Likewise, the secondary and tertiary structure (**Fig. S4c**), resistance to chemical and thermal denaturation (**Fig. 5c, S4d**) and specific activity (**Fig. 5d**) of purified CAT produced from the Shuf1 mRNA sequence were indistinguishable from CAT translated from the WT coding sequence. We also used mass spectrometry to compare the molecular weights of CAT translated from these coding sequences. These masses were indistinguishable to within one mass unit and matched the expected molecular weight of 25,953 Da. Taken together, these results demonstrate that CAT production from the Shuf1 coding sequence does not prohibit formation of a stable, active CAT protein structure.

Despite the native CAT structural similarities reported above, it is important to note that digestion by ClpXP requires force-mediated unfolding of a substrate protein from its C-terminus, driven by ATP hydrolysis (51–53). Resistance to mechanical force reports on a distinct aspect of protein stability from resistance to chemical or thermal denaturation (54–57). To directly test whether the Shuf1 synonymous codon substitutions lead to a native CAT structure that is more susceptible to force-mediated unfolding and degradation, we subjected native, purified ssrA-tagged CAT produced *in vivo* from the WT or Shuf1 coding sequences to an *in vitro* ClpXP degradation assay (44, 58). Although both proteins exhibit resistance to ClpXP degradation, CAT synthesized from the Shuf1 coding sequence was degraded more rapidly than WT-CAT (**Fig. 5e, S5**), in direct contrast to the indistinguishable behavior observed in our other analyses (above). This result demonstrates that CAT produced from the Shuf1 mRNA sequence is more susceptible to degradation by ClpXP, perhaps both prior to as well as after acquiring its native structure. Crucially, the differential susceptibility to ClpXP degradation provides direct evidence of the impact of the Shuf1 codon substitutions on CAT folding, as proteins with identical amino acid sequences would arrive at different native structures only via distinct folding mechanisms. Because the ssrA tag is located at the CAT C-terminus, we expect that degradation by ClpXP is predominantly post-translational, occurring after release of the nascent chain from the ribosome.

### mRNA secondary structural stability does not explain Shuf1 growth defect

The results above suggest the Shuf1 synonymous codon substitutions impair CAT co-translational folding by altering local patterns of translation elongation. *In vitro*, synonymous codons have been shown to alter elongation rate either by altering the rate of decoding (59) or by altering downstream mRNA stability, which can impede ribosome translocation (60). *In vivo*, there is some evidence that stable mRNA stem-loop structures can alter the elongation rate of the ribosome (61–63), although other studies have detected no difference (38, 64, 65), likely due to destabilization of mRNA structure by polysomes and/or the helicase activity of the ribosome. Although the overall predicted mRNA stability of the WT and Shuf1 genes are similar, a predicted stable 3’ stem-loop structure in Shuf1 is not present in the WT coding sequence (**Fig. S6a**). To test whether this structure is responsible for the Shuf1 growth defect, we created chimeric mRNA sequences with only the 5’, middle or 3’ portion of the wild type sequence substituted with the Shuf1 sequence (**Fig. S6b**) but observed no growth defect for the chimera bearing the 3’ portion of Shuf1 had no impact on growth rate (**Fig. S6c**). Moreover, growth rates for these chimeras correlated more closely with the difference in relative codon usage frequencies than measures of mRNA stability (**Fig. S6d**). Taken together, these results indicate that translation elongation rate differences arising from changes in codon usage frequencies is a more likely origin of the Shuf1 growth defect than changes in mRNA secondary structure.

## Discussion

Most of our current understanding of protein folding mechanisms is derived from studies of small proteins that refold reversibly when diluted from chemical denaturants. However, only a small number of proteins can refold robustly *in vitro*, even though many more can be maintained in a stable state once extracted from the cell (24, 66, 67). This suggests that (i) the conformations adopted early during the folding process are crucial to successful folding and (ii) the cellular environment supports the formation of early folding intermediates that are distinct from the conformations populated upon dilution from denaturant. Indeed, there is substantial evidence that molecular chaperones are crucial to the successful folding of many complex proteins *in vivo* (29–33). Although it has been hypothesized that synonymous codon changes could alter elongation rate and modify folding mechanisms *in vivo*, it has thus far been challenging to find evidence to support this hypothesis from experiments performed *in vivo*, possibly due to buffering provided by molecular chaperones.

Results presented here indicate that, during synthesis, the folding of nascent CAT polypeptide chains is sensitive to synonymous codon-induced changes to translation elongation rate. Although the nascent chains produced using different synonymous codon patterns remain broadly capable of achieving a stable, active CAT trimer structure, translation using the synonymous Shuf1 mRNA sequence leads to CAT proteins that are more susceptible to degradation by the cellular protease ClpXP than WT-CAT, leading to a dramatic cell growth defect. Given that the ClpXP ssrA degradation tag is attached to the very C-terminus of CAT, it is likely that the majority of this digestion occurs only post-translationally, after the CAT nascent chain is released from the ribosome (**Fig. 4**). Remarkably, even native Shuf1-CATssrA protein is more susceptible to degradation than native CATssrA translated using the WT coding sequence, demonstrating that the impact of the codon-induced perturbations persists long after translation and folding is complete. Buffering by the cellular protein homeostasis network is therefore not sufficient to mask the impact of the Shuf1-CAT folding defect on cell growth.

These results are consistent with a small but growing number of studies indicating that synonymous codon substitutions can perturb protein folding mechanisms (1, 10, 12, 68). The ssrA tagging approach developed here provides a general strategy to uncover such perturbations in other coding sequences, even when they do not lead to dramatic remodeling of the final protein structure. In contrast to the translation rate-sensitive effects we observed for CAT folding, recent *in vitro* single molecule force-unfolding experiments have shown that some small, ribosome-bound natively-folded domains can fold via similar mechanisms on and off the ribosome (69, 70). However, as these studies noted, forced unfolding measured by molecular tweezers cannot capture the transient folding of a nascent chain during its synthesis (33), and hence what is measured in these experiments is the effect of close proximity of the ribosome surface, rather than co-translational folding. The very robust folding behavior of these well-characterized, reversible folding models may indeed lead to indistinguishable folding behavior during translation, a model supported by recent force-feedback folding measurements (71). However, the model proteins selected for these studies are smaller than >75% of proteins in the *E. coli* proteome (24), whereas all known examples of synonymous codon-derived alterations to co-translational folding are much larger (e.g., (1, 9, 10, 72)). We are not aware of an *in vitro* folding mechanism for a protein >175 aa long that is preserved during co-translational folding. Synonymous codon-derived modulation of elongation rate may therefore play a broad role in the efficient folding of larger, more complex proteins.

Our CAT results demonstrate that synonymous changes to mRNA coding sequences can significantly perturb folding of the wild type protein sequence even in the presence of the cellular repertoire of molecular chaperones. This result suggests that mRNA sequences have likely evolved alongside molecular chaperones to most efficiently support folding of the broad repertoire of protein structures produced *in vivo*. Although our understanding of co-translational folding mechanisms is still in its infancy, these results imply that it should ultimately be possible to rationally design mRNA coding sequences to enhance *in vivo* folding yield and to identify disease-associated synonymous codon substitutions most likely to adversely affect protein co-translational folding, particularly for large or otherwise complex proteins.

## Methods

### Cell growth assays

A single colony of *E. coli* KA12 (73) or W3110 (50) transformed with a pKT-CAT plasmid from a freshly streaked LB-amp plate was used to inoculate 20 mL of LB plus 100 μg/mL ampicillin (LB-amp) and grown overnight with shaking at 37°C. Unless otherwise specified, all cultures contained 100 μg/mL ampicillin and no tetracycline. Overnight cultures were used to innoculate fresh LB-amp to an optical density at 600 nm (OD_600_) of 0.05, to which was added 35 μg/mL chloramphenicol (unless otherwise specified) and the indicated concentration of tetracycline inducer (0-1600 ng/mL), transferred to one well of a 12-well plate and incubated at 37°C with continuous shaking in a Synergy H1 microplate reader (BioTek). Growth was measured as the increase in OD_600_. The linear portion of the growth curve was fit to a straight line and the slope was taken as the growth rate.

## Acknowledgements

We thank Matt Champion and the Notre Dame Mass Spectrometry & Proteomics Facility for performing the mass spectrometry experiments, Don Hilvert for the kind gift of the pKT and pKTS plasmids and Peter Chein, Don Hilvert, Jeff Nivala and Mark Akeson for sharing *E. coli* strains with us. We are grateful to Gabriel Wright and Scott Emrich for helpful discussions. This project was supported by grants GM120733 and GM105816 from the National Institutes of Health.

## Supplemental Information

### Supplemental Methods

#### Synonymous CAT coding sequence design

The Shuf1 synonymous mutant sequence was designed using a Monte Carlo algorithm. The first 46 codons (excluding the His tag) were left unchanged to avoid altering translation initiation (5, 38–41). For remaining amino acid positions, a synonymous codon was chosen at random. After each random sequence was designed, its codon usage frequencies were analyzed using the %MinMax algorithm (18, 37), which compares the codon usage frequency of a coding sequence to hypothetical sequences that encode the identical amino acid sequence using either the most common (+100%) or most rare (−100%) codons possible. This simple algorithm has been shown to accurately predict the relative effects of synonymous codon changes on elongation rate (10, 19). The %MinMax codon usage profile of each synonymous sequence was compared to the WT profile. Profiles were filtered to select those with the same number of 17-codon windows of rarer-than-average codon usage, a sequence-wide average codon usage very similar to the wild type mRNA sequence (%MinMax within +/− 3% of WT), and likewise very similar extreme codon usage values (maximum and minimum %MinMax values within +/− 5% of WT). Out of one million sequences generated, three fulfilled these criteria. The coding sequence with the lowest Pearson correlation coefficient compared to WT was chosen as Shuf1.

#### Molecular biology

All CAT constructs were synthesized by GeneArt (Invitrogen). The pKT and pKTS plasmids were a generous gift from Donald Hilvert (ETH, Zurich). CAT constructs were cloned into pKT and pKTS using FastCloning (74). All primers were ordered from IDT as standard desalting primers and resuspended to 1 mg/mL in ddH_2_O. pKT and pKTS were amplified using primers pKT F (ACTAGTGCGGCCGCTTGATAAA) and pKT R (ATGTATATCTCCTTCTTAAAGTTAAACAAAATTATTTCTAGAGGGAAA) and pKTS F (GCGGCGAACGATGAAAACTATGC) and pKT R, respectively. The CAT F primer (AGAAGGAGATATACATATGCATCACC ATCACCATCACCATAACTATACAAAATTTG) was used to amplify all CAT constructs. Unique primers were required for the reverse direction. WT R pKT (AAGCGGCCGCACTAGTTTATTTTAATTTACTGTTACACAACTCTTGTAG) was used to clone WT and the common variant of CAT (11) into pKT, WT R pKTS (TTTCA TCGTTCGCCGCTTTTAATTTACTGTTACACAACTCTTGTAGCCGATTAATAAAGC) for pKTS, Shuf1 R pKT (AAGCGGCCGCACTAGTTTACTTCAATTTAGAATTACATAACTCCTGTAAT) was used to clone Shuf1 CAT into pKT, and Shuf1 R pKTS (TTTCATCGTTCGCCGCCTTCAATTTAGAATTACATAACTCCTGTAATCTGTTAATGAAGC) for pKTS.

To make the chimeric CAT constructs, the 5’, center or 3’ region of the Shuf1 coding sequence were amplified and cloned into the WT CAT sequence. All hybrid CAT constructs were cloned into pKTS. Primers were ordered from IDT as standard desalting primers. To make the 5’ chimera, the WT plasmid was amplified using the primers WTsN F (GCTATCCTGCCCC TATTCATCCGATATTGATCAATTTATGGTGAATTATTTATCGGT) and WTsN R (ATATCATCA CGGGGTAAAACTTATACGCTGAATCATCCAATGACTTTTTTAACGTC) and codons 58-111 of Shuf1 using the primers Shuf1 N F (TACCCCGTGATGATATACTTAATTGCCCA) and Shuf1 N R (ATAGGGGCAGGATAGCGCAC). To make the middle chimera, the WT plasmid was amplified using the primers WTsM F (TGACTATTTTGCGCCGATTATAACAATGGCAAAATATCAGCAAGAAGGG) and WTsM R (GATCGATGTCCGAGCTGTATGGGCAACTCAGTGCTGAAAA) and codons 112-172 of Shuf1 using the primers Shuf1 M F (AGCTCGGACATCGATCAATTTATGG) and Shuf1 M R (CGGCGCAAAATAGTCAGTAAAGTTAGCTA). To make the 3’ chimera, the WT plasmid was amplified using the primers WTsC F (TAATTCTAAATTGAAGGCGGCGAACGATGAAAACTATGC) and WTsC R (TGGCCATGGTTATGATGGGTGCAAAATAATCGGTAAAATTAGCAA) and codons 173-220 of Shuf1 using the primers Shuf1 C F (ATCAT AACCATGGCCAAATACCAACAG) and Shuf1 C R (CTTCAATTTAGAATTACATAACTCCTGTAATCTGTTAATGAAGC).

PCR reactions were prepared using 5 μL of supplied 10x *Pfu*Ultra II buffer (Agilent), 2.5 mM dNTPs (AMRESCO), 2 ng/μL of forward and reverse primer, 0-3 mM MgCl_2_, 0.2 ng/μL DNA, and 1 μL *Pfu*Ultra II Fusion HS DNA polymerase (Agilent). The total reaction volume was 50 μL. The PCR protocol started with 2 min at 95°C followed by 18 cycles of: 95°C for 30 sec, T_m_-5°C for 1 min, 72°C for 2.5 min, then a final step of 72°C for 10 min.

PCR products were treated with 1 μL *DpnI* for 1.5 hours at 37°C. The insert and vector were then mixed together in 1:1, 1:2, 1:4, 2:1, and 4:1 ratios and 10 μL were incubated with 100 μL chemically competent *E. coli* DH5α for 30 min on ice. Cells were heat shocked at 42°C for 45 sec, incubated on ice for 2 min, and recovered at 37°C with 500 μL of SOC media (2% tryptone, 0.5% yeast extract, 10 mM NaCl, 2.5 mM KCl, 10 mM MgCl_2_, 10 mM MgSO_4_, 20 mM glucose) for 1 hr. The entire transformation reaction was spread on an LB-agar plate with 100 μg/mL ampicillin and incubated overnight at 37°C. Single colonies were grown in liquid culture and DNA was extracted (Wizard Plus minipreps, Promega) using the manufacturer’s directions. The entire gene along with its promoter and transcription terminator was sequenced.

#### CAT protein purification

To purify CAT for *in vitro* analyses, 1 L cultures were inoculated as above with 1600 ng/mL tetracycline. Cultures were grown at 37°C with continuous shaking for 6.5 hours. Cells were pelleted at 4,000xg for 15 min and resuspended in nickel affinity binding buffer (500 mM NaCl, 20 mM imidazole, 20 mM sodium phosphate pH 8). Cells were lysed by repeated cycles of freezing at −80°C for 30 min followed by thawing at room temperature. Lysozyme (final concentration 1 mg/mL) was added after the first freezing cycle. Thawed lysates were incubated for 30 min at 4°C with frequent inversions to facilitate lysis. In total, cells were frozen and thawed four times. After lysis, cells were homogenized with 25 mM MgCl_2_ and 80 μL DNase I (Invitrogen). The lysates were again incubated at 4°C for 30 min with frequent inversions. Alternatively, lysates were homogenized by sonication for 12 min using 30 sec on 30 sec off. After lysis, the lysate was pelleted at 14,000xg for 15 min and CAT was purified from the supernatant using nickel affinity chromatography (His Spin Trap (small volumes), HisTrap HP (large volumes); GE Healthcare). After addition of the cell supernatant, columns were washed with binding buffer and eluted with elution buffer (500 mM NaCl, 500 mM imidazole, 20 mM sodium phosphate pH 8). CAT protein concentration was determined by absorbance at 280 nm, using a calculated CAT extinction coefficient of 30,350 (for His-CAT) or 31,630 M^-1^ cm^-1^ (for His-CAT-ssrA) (75).

#### Far-UV circular dichroism spectroscopy

Far-UV CD spectra were collected using a J-815 CD Spectropolarimeter (JASCO). CAT protein samples were diluted to a concentration of 4 μM in 20 mM phosphate buffer pH 8. Three spectra were collected from 190-250 nm with a 1 second integration time and a 2 nm bandwidth, averaged and from this an averaged spectrum of the blank buffer solution (collected under identical conditions) was subtracted. The effect of increasing temperature on the far-UV CD spectrum was recorded every 2°C from 4-95°C. Samples were equilibrated for 2 min at each temperature prior to data collection. To normalize the spectra, the average signal prior to and after the unfolding transition were set to 100% and 0% folded, respectively.

#### Tryptophan fluorescence emission spectroscopy

Purified CAT protein was diluted to 2 μM and incubated overnight in 100 mM Tris buffer pH 7.5 with 0-4 M guanidinium HCl at 4°C. Tryptophan emission spectra were collected using a PTI QM-6 fluorimeter (Birmingham, NJ). An excitation wavelength of 280 nm was used (2 nm slit widths) and emission was measured from 300-380 nm (4 nm slit widths; 3 sec integration time). To normalize the spectra, the average fluorescence emission intensity prior to and after the unfolding transition were set to 100% and 0% folded, respectively. The resulting curves were then fit to a sigmoidal function.

#### Acetyltransferase activity assay

CAT enzymatic activity was measured as described previously (76). Briefly, CAT was diluted to a concentration of 6.25 μg/mL in 1 mL of reaction mix (94 mM Tris pH 7.8, 83 μM DTNB, 190 μM acetyl-CoA and 155 μM chloramphenicol). Free CoA-SH produced by the CAT reaction subsequently reacts with DTNB to make TNB, which absorbs at 412 nm. Absorbance at 412 nm was monitored for 5 min in a DU 530 UV-Vis Spectrophotometer (Beckman Coulter). To determine the specific activity from these curves, the initial velocity of the reaction was calculated by fitting data points from 0.25-1 min to a straight line. The initial velocity was divided by the extinction coefficient for TNB (0.0136 μM^-1^ cm^-1^) to convert to units of enzyme activity and the amount of CAT to calculate activity in units per microgram. One unit of activity converts 1 nmol of chloramphenicol and acetyl-CoA to chloramphenicol-3-acetate and CoA per min. The average activity of WT His-CAT was set to 100%; results for other CAT proteins were normalized to this value.

#### Mass spectrometry

The molecular weight of CAT was measured using HPLC-ESI-MS as described previously (77, 78), with minor modifications. Briefly, purified CAT was diluted to 0.5 mg/mL in 25 mM ammonium bicarbonate pH 8.0 and 5 μg was injected per sample. A 20 min gradient from 10-80% A-B (A: H2O, 0.1% formic acid; B: acetonitrile, 0.1% formic acid) with the first 3 min diverted to waste was performed. Positive mode electron spray ionization was acquired on a micro TOF-QII (Bruker, Billerica, MA) mass spectrometer with 2 sec spectral averaging. The MaxEnt software package (Bruker) was used for deconvolution. The deconvoluted mass measured for each CAT variant was 25,953 Da, identical to the predicted mass.

*E. coli* expressing WT or Shuf1 His-CAT-ssrA were grown to an OD_600_ of 0.3 at 37°C in LB with amp and induced with 1000 ng/mL tc for 3 hr. Cells (from 10 mL cultures) were pelleted at 4,000xg for 15 min and stored at −80°C. Cell pellets were lysed using a bead mill (BioSpec OK, USA) 3 x 30 sec in 1% SDS 100 mM ammonium bicarbonate (ABC) and clarified by centrifugation at 12,000xg for 10 min. Whole cell protein lysates were quantified by BCA assay (Thermo Scientific) according to the manufacturer’s instructions.

Whole cell lysates were prepared for proteomics as previously described (79), with minor modifications. All reagents were from Sigma unless otherwise described. First, 50 μg of each lysate was suspended in 5% SDS 100mM ABC and reduced and alkylated with dithiothreitol and iodoacetamide. Samples were then digested with trypsin using Strap columns (Protifi, NY) according to manufacturer’s instructions, with the following alterations. Mass-spectrometry grade trypsin (Promega, WI) was added at 1:40 (m/m) and digested overnight at 37°C. Peptides were eluted from the digestion column and desalted using a 10 mg HLB SPE column (Waters, MA) following the manufacturer’s recommended conditions. Desalted peptide digests were dried and stored at −20°C until LC/MS/MS analysis.

Each digested sample (1 μg) was analyzed in triplicate using LC-MS/MS with a Waters nanoUHPLC Acquity (Billerica, MA) and a Q-Exactive Oribitrap (Thermo, San Jose, CA). RAW files from the Nano UHPLC-MS/MS acquisition were subjected to tandem spectral match and quantification using label free quantification (80). The current KA12 *E. coli* proteome (UP000000625) concatenated with the WT/Shuf1 His-CAT-ssrA sequences and common contaminants was used to assign peptides. Data were normalized using median fold change.

#### Calculation of local mRNA secondary structure

A sliding window analysis was used to predict local stable structure in overlapping 30 nt-long segments of the CAT coding sequence, using the Quikfold algorithm to calculate the free energy of mRNA folding (81). All predicted stable structures were plotted.

#### Correlation between growth and mRNA properties

Growth was calculated as the %WT OD_600_ after 4 hours of growth in cam. The change in %MinMax was calculated by summing the absolute value of the differences between a given construct and WT. This number was divided by the average %MinMax of the gene as a proxy for protein production. The strongest local mRNA secondary structure element per gene was calculated as above. These data were fit to a line using Prism.

#### Western blotting

Cultures were inoculated as described above and induced for 6.5 hours. Aliquots were pelleted at 14,000xg for 15 min and the cell pellets were resuspended to an OD_600_ of 15 in 100 mM Tris pH 7.5. To this was added one half-volume of 3x SDS gel loading buffer and β-mercaptoethanol (BME) to 5 mM. Samples were boiled for 40 min, separated using denaturing polyacrylamide gel electrophoresis, transferred to a PVDF membrane and probed using a mouse anti-His antibody (Promega) and goat anti-mouse secondary antibody conjugated to alkaline phosphatase. In some experiments, the 7x-His tag was replaced with an HA tag (YPYDVPDYA) and the primary antibody used was rabbit α-HA (Sigma). Intensities of CAT protein bands were quantified using ImageJ.

#### mRNA quantification

Cultures were inoculated as described above and grown for 2.5 hr until the cells achieved an OD_600_ of about 0.3. RNA was purified using the RNeasy kit (Qiagen), according to manufacturer protocol. RNA concentrations were determined by absorbance at 260 nm. Purified RNA was stored at −80°C.

RNA samples were treated with DNase to remove genomic DNA or plasmid contaminants. To 10 μL of RNA (about 0.2 μg) 10 μL of DNase buffer (10 mM Tris pH 7.5, 2.5 mM MgCl2, 0.5 mM CaCl_2_) was added with 2 units of RNase-free DNase (Ambion/Invitrogen). Samples were incubated at 37°C for 30 min, then 1 μL of 100 mM EDTA was added and samples were incubated at 65°C for 10 min to denature the DNase. Reverse transcription (RT) was performed using the iScript RT kit (BioRad). RNA was diluted to about 100 ng/μL into RT reaction mix (20 μL total, 4 μL 5x buffer, 2 μL random hexamer primers, 1 μL reverse transcriptase). Control reactions lacking RT were used to ensure the PCR amplification was specific for mRNA. To perform the RT, the following protocol was used on a MyCycler (BioRad): 5 min at 25°C, 30 min at 42°C, 5 min at 85°C. The resulting cDNA was diluted fivefold for quantitative PCR (qPCR) in a 96 well plate using a CFX96 Touch Real-Time PCR Detection System (BioRad). The reaction volume was 5 μL and included 1 μL of cDNA, 60 nM primers (IDT) and 2.5 μL SsoAdvanced SYBR Green Supermix (BioRad). To amplify all CAT constructs the primers CAT qPCR F (CCATCACCATAACTATACAAAATTTGATGTAAAAAATT) and CAT qPCR R (AAACTTATACGCTGAATCATCCAATGA) were used. To amplify *E. coli* GAPDH, the primers GAPDH qPCR F (CGGTACCGTTGAAGTGAAAGAC) and GAPDH qPCR R (ACCAGTTGCTTCAGCGAC) were used. ΔC_T_ values were calculated by taking the difference in C_T_ values for each CAT construct with its corresponding GAPDH C_T_. ΔΔC_T_ values were calculated by taking the difference in ΔC_T_ from WT CAT. This was converted to fold change in RNA level by raising 2 to the power of -ΔΔC_T_.

#### Growth rate as function of induction time

Cultures were inoculated as described above in 20 mL of LB with amp and grown in shake flasks at 37°C. Cultures were grown to an OD_600_ of ~0.35 at 37°C and induced with 1600 ng/mL tc. Cells were induced from 0-60 min. Then a 1 mL aliquot was pelleted at 21,000xg for 5 min and resuspended in LB with amp and cam. These were diluted twofold into a 24 well plate. Growth was monitored with double orbital shaking for 5 hours. Growth from 1-3 hours after chloramphenicol addition was fit to a line and the slope was used as the growth rate. Growth rates were normalized to the average growth rate of both constructs after 60 min induction.

#### ClpX knockout effect on growth

The ClpX knockout was in the *E. coli* W3110 background. Knockout and parent strain were cultured as described above. These strains were induced with 50 ng/mL tc. The effect of ClpX or ssrA tag on growth was calculated by dividing the endpoint OD_600_. No effect of deletion would render a value of 1.

#### Contact maps

Contact maps were generated using VMD software support and the crystal structure of CATIII (PDBID: 3CLA) (34). VMD was developed with by the Theoretical and Computational Biophysics group at the Beckman Institute, University of Illinois at Urbana-Champaign.

#### ClpXP degradation assay

ClpXΔN_3_ or ClpP were overexpressed in *E. coli* and purified using Ni^2+^ affinity and size exclusion chromatography, as described previously (82). ClpXP degradation were performed in PD buffer (25 mM HEPES pH 7.6, 100 mM KCl, 10 mM MgCl_2_, 10% v/v glycerol, 2 mM BME at 30°C. CATssrA purified as above (2 μM) was incubated with 300 nM ClpX_6_ (as 600 nM ClpXΔN_3_), 900 nM ClpP_14_, 5 mM ATP (Sigma), 0.2 mg/ml creatine phosphokinase (CPK, Sigma), and 1.6 mM creatine phosphate (Sigma). At indicated timepoints, an aliquot of the degradation reaction mix was removed and combined with SDS-PAGE loading dye. The degradation reaction was monitored by SDS-PAGE.

**Figure S1.**
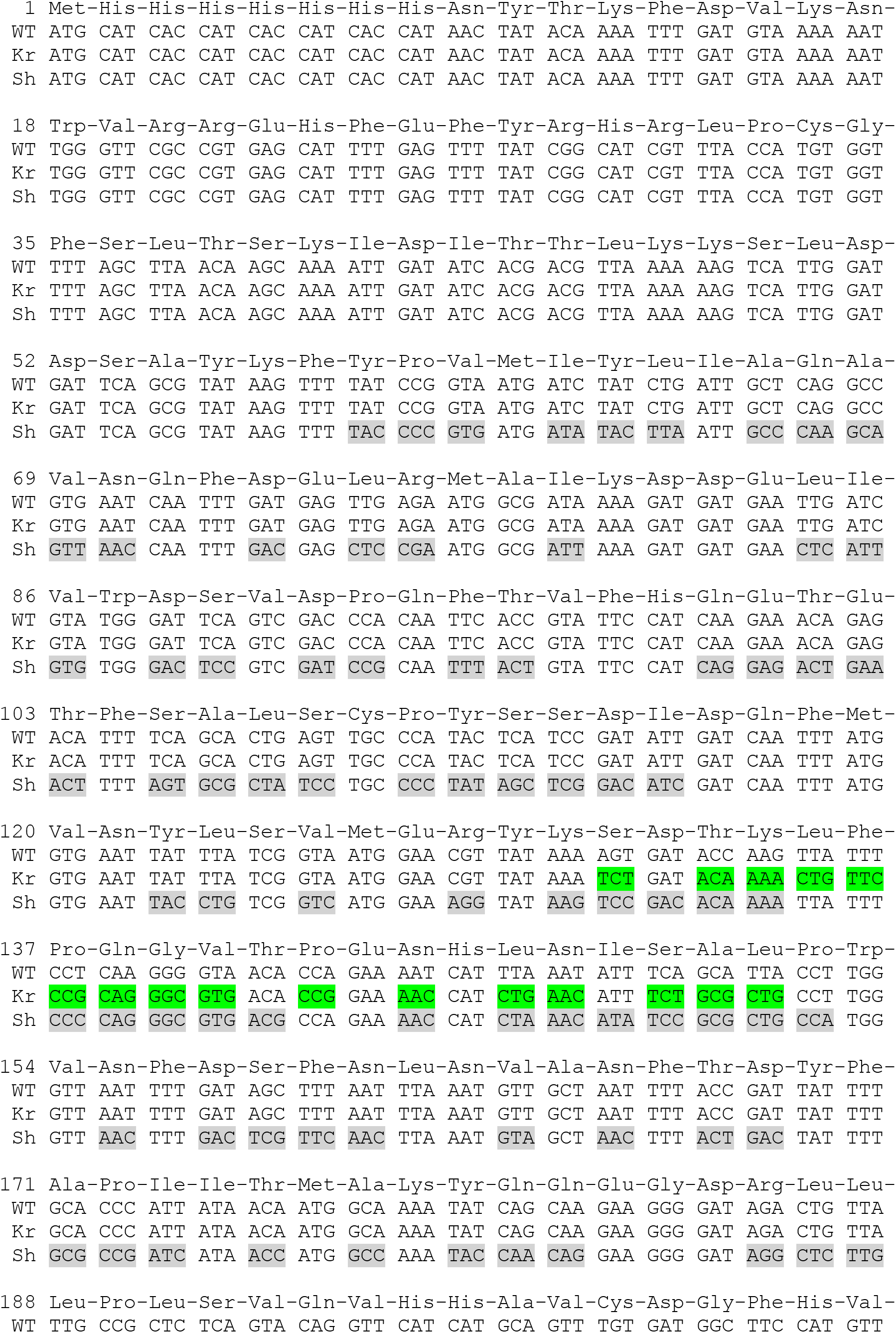
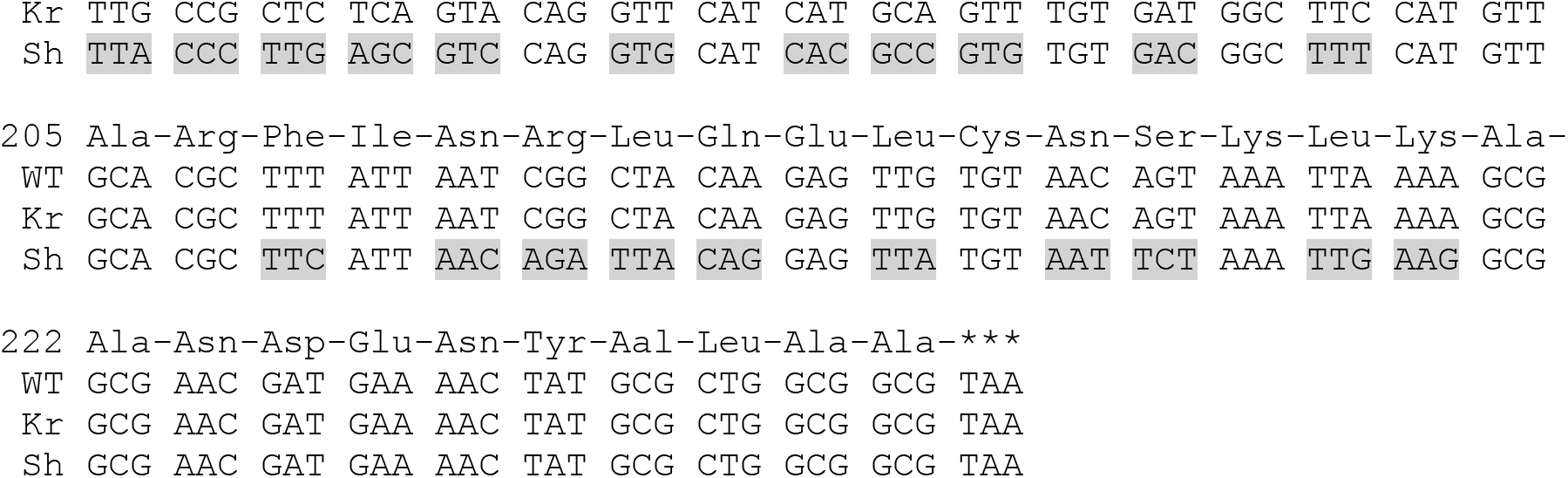
CAT coding sequences used in this study. Codon substitutions are highlighted. WT, wild type CAT; Kr, synonymous variant described previously [11]; Sh, Shuf1 synonymous variant.

**Figure S2.**
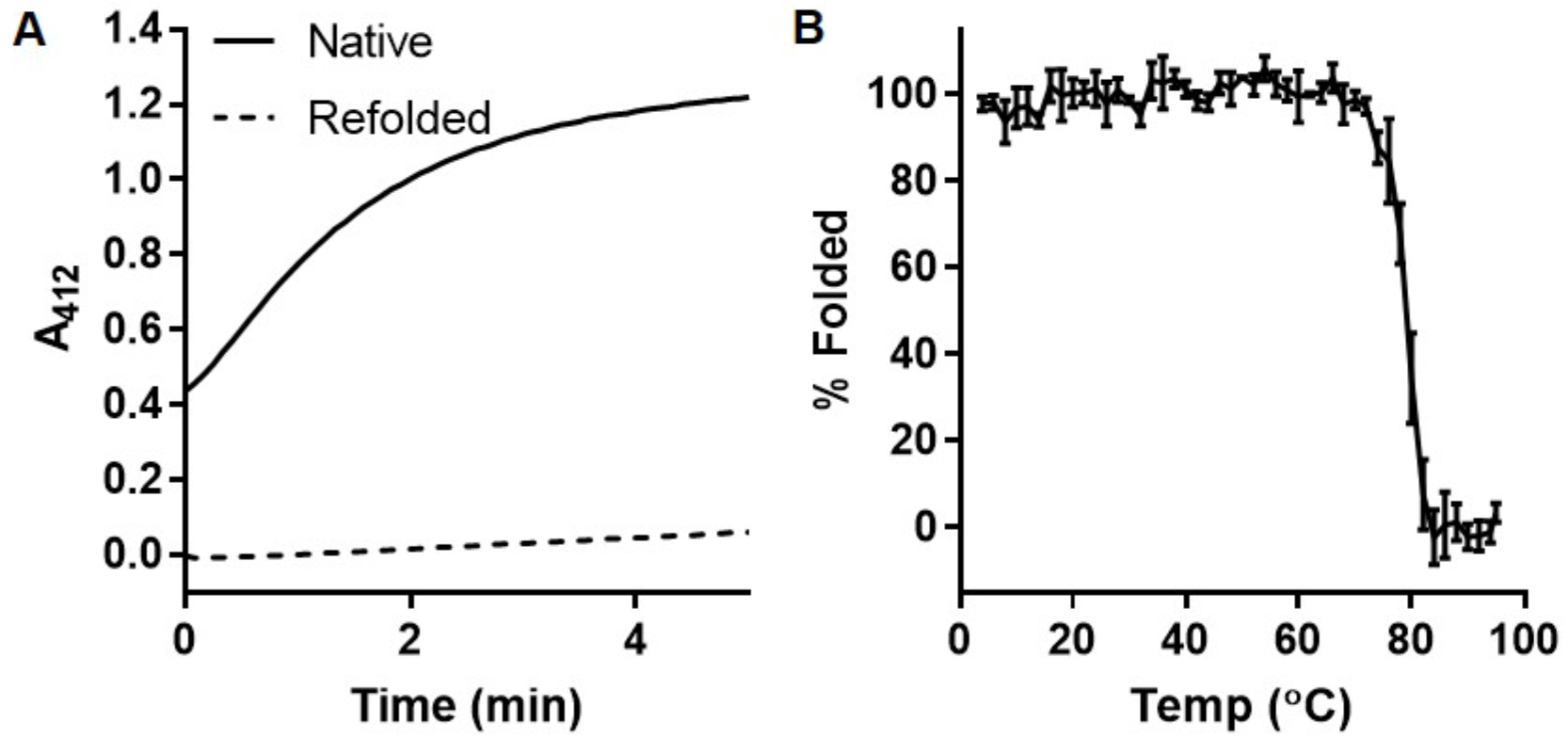
The native CAT structure is very thermostable but cannot be refolded after chemical denaturation. (**A**) CAT activity, expressed as the change in absorbance of TNB over time in the presence of native or refolded CAT (see Methods). Refolded CAT does not have significant acetyltransferase activity. (**B**) Thermal denaturation of native, purified CAT, monitored as the change in the far-UV CD signal at 205 nm; n=3, data are represented as mean ± SD.

**Figure S3.**
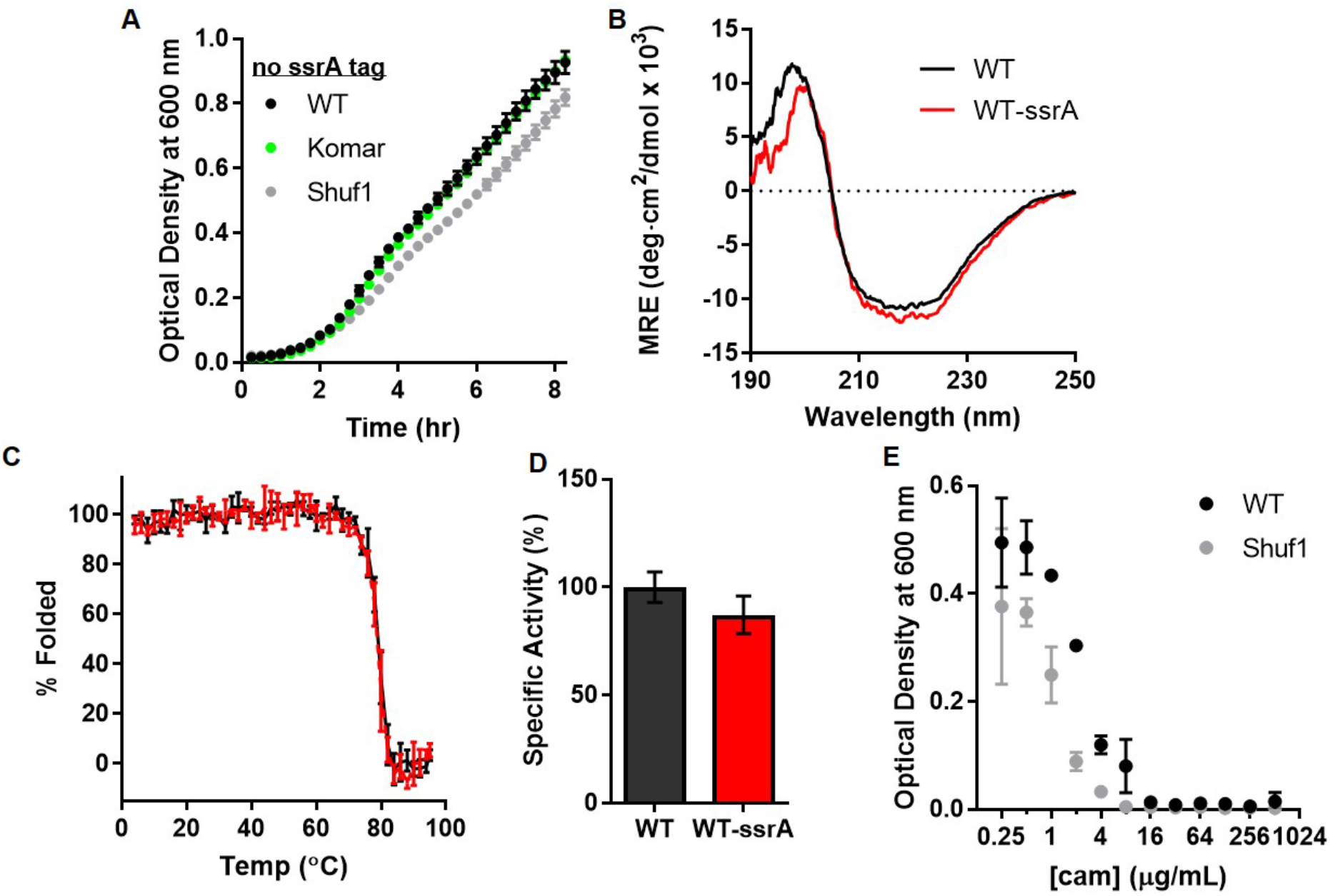
The C-terminal ssrA tag does not affect properties of native, purified CAT, but does affect chloramphenicol resistance of *E. coli* expressing CAT from the Shuf1 coding sequence. (**A**) Growth curves for cell expressing CAT codon variants without a C-terminal ssrA tag. (**B**) Far-UV CD spectra of WT CAT and CAT-ssrA. (**C**) Thermal denaturation of WT CAT and CAT-ssrA monitored by changes in the far-UV CD signal at 205 nm. (**D**) Acetyltransferase activity of WT CAT and CAT-ssrA are indistinguishable; p=0.12, Welch’s t-test; n=3, data are represented as mean ± SD. (**E**) Minimum inhibitory concentration of chloramphenicol (cam) for *E. coli* cells expressing WT- or Shuf1-CATssrA.

**Figure S4.**
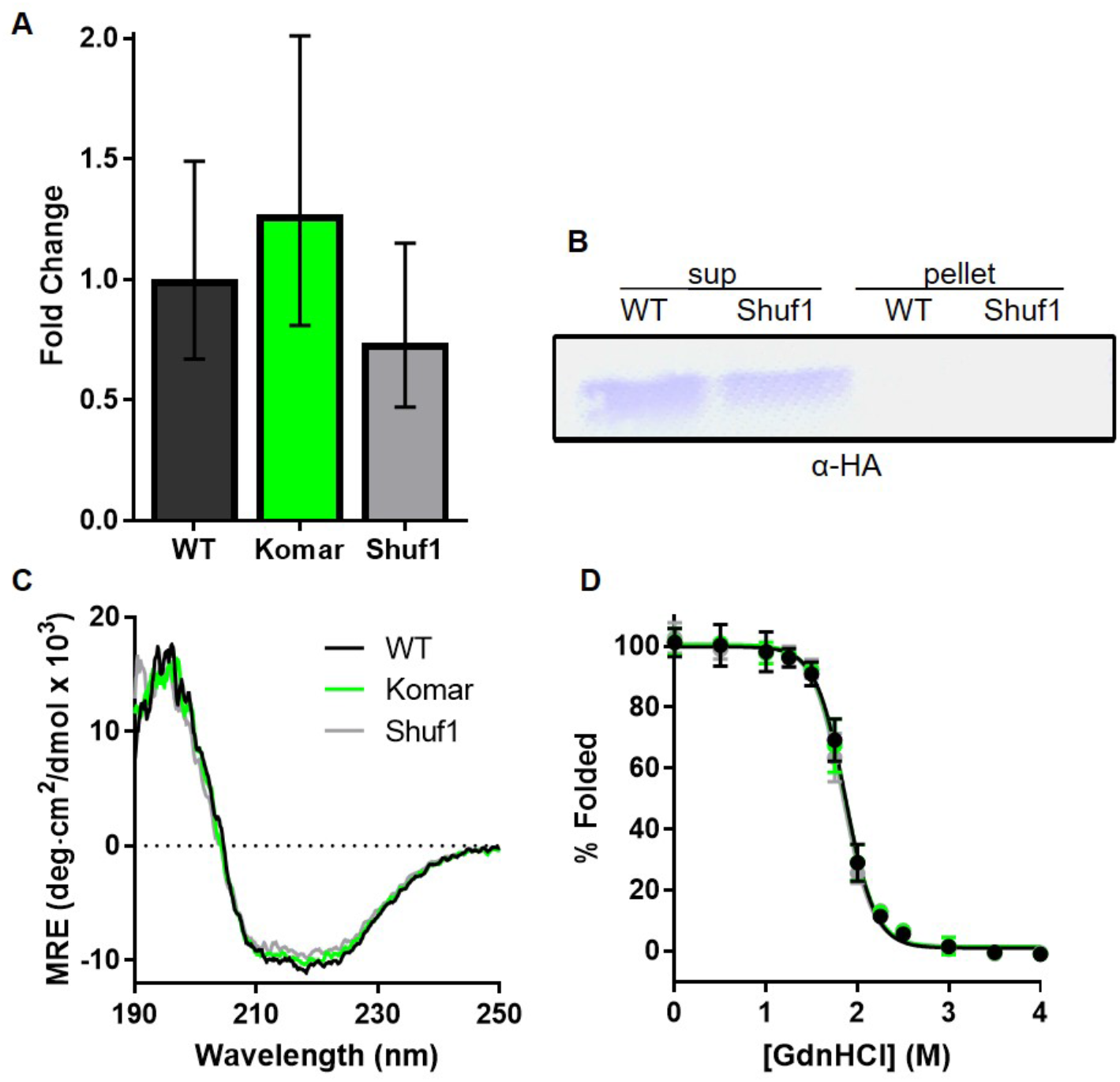
CAT mRNA concentration and protein solubility, secondary structure and resistance to chemical denaturation are indistinguishable, regardless of coding sequence. (**A**) Fold change in CAT mRNA levels relative to WT, as determined by RT-qPCR. (**B**) Partitioning of CAT-ssrA into soluble (sup) and insoluble (pellet) cell lysate fractions. (**C**) Far-UV CD spectra. (**D**) Guanidinium hydrochloride denaturation of CAT monitored as change in ratio of tryptophan fluorescence emission intensity at 330 versus 349 nm; n=3, data are represented as mean ± SD.

**Figure S5.**
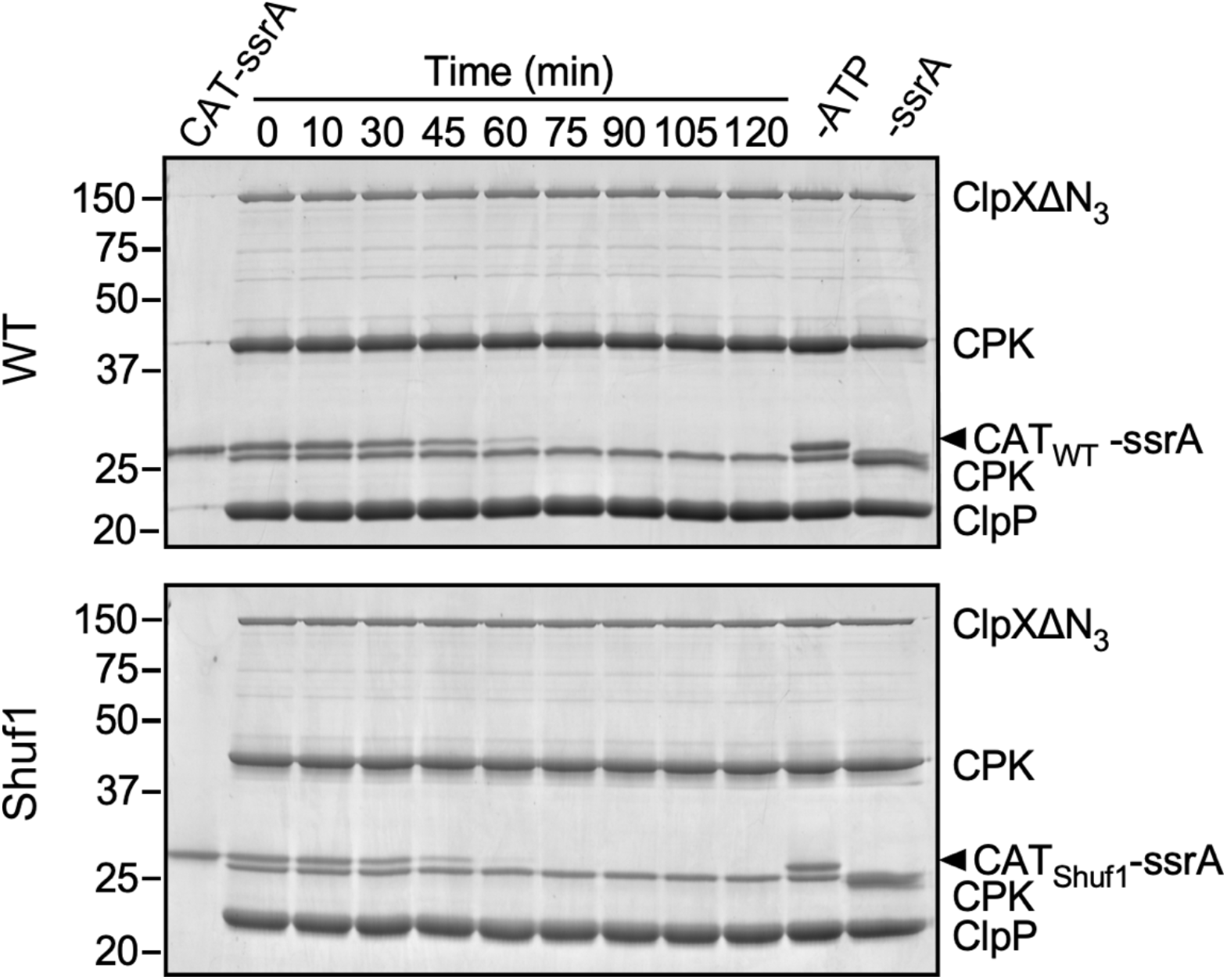
One biological replicate of SDS-PAGE gel results from ClpXP degradation assay, quantified in Figure 5e. Bands marked CPK correspond to creatine phosphokinase from the ATP regeneration mixture (see Methods).

**Figure S6.**
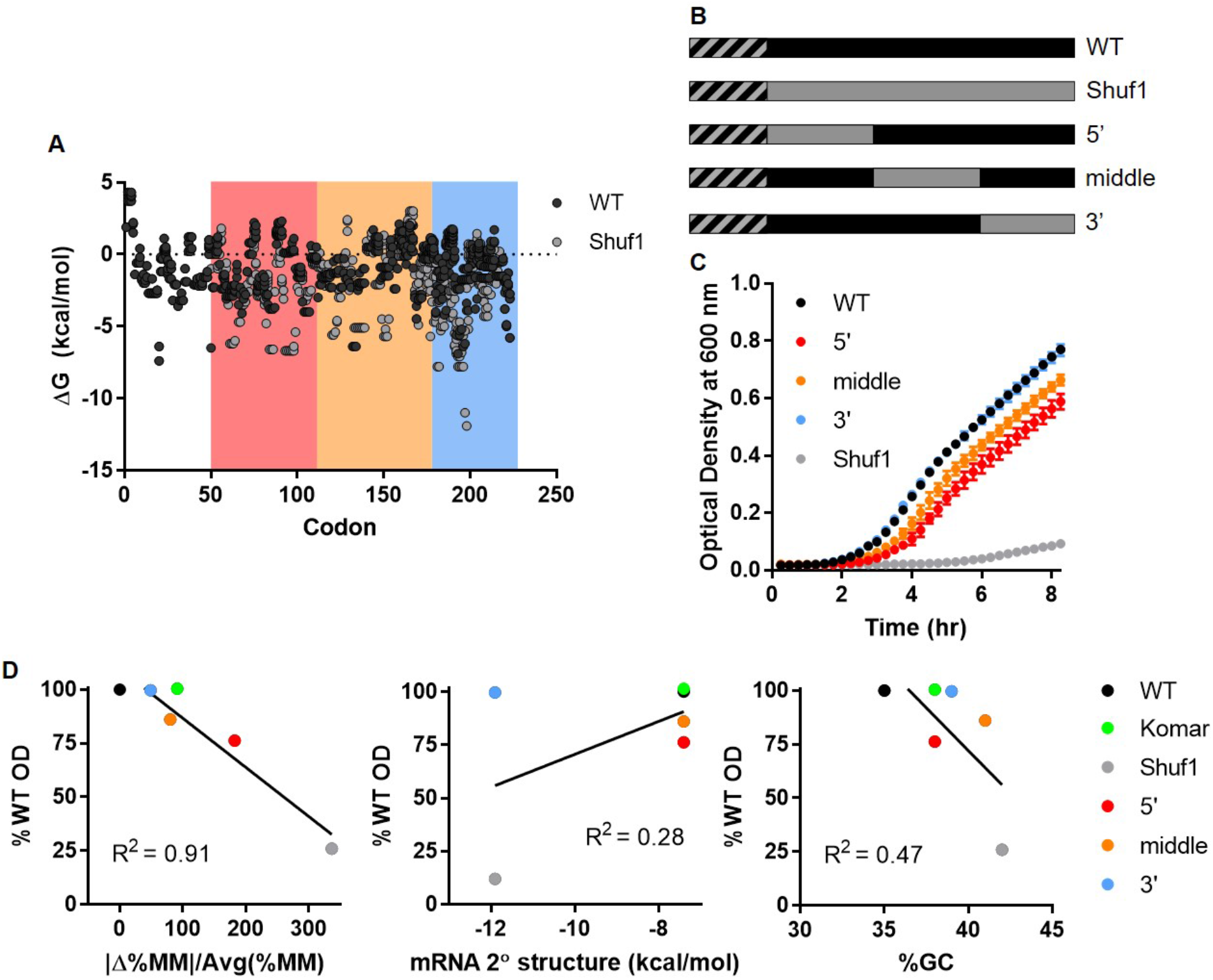
Growth of *E. coli* expressing chimeric CAT mRNA coding sequences. (**A**) mFold predictions of local mRNA secondary structure over 30 nt windows. Background color indicate chimera construct boundaries. (**B**) Schematic of chimeric mRNA sequences; black, WT; gray, Shuf1. The hatched segment represents the most 5’ 46 codons, which were identical in each sequence. (**C**) Growth curves for the 5’, middle, and 3’ chimeras compared to WT and Shuf1. Data points represent the mean ± SD; n=3 biological replicates. (**D**) Correlations between *E. coil* growth and various CAT chimeric coding sequence properties, including (left) difference in codon usage frequency (plotted as the change in %MinMax divided by the average %MinMax), (middle) strongest local mRNA secondary structure element, and (right) %GC content. Growth is plotted as the percentage of the WT OD_600_ after 8 hr of growth. Growth correlates most closely with codon usage.

## Notes

#### Summary of Updates

We repeated the in vitro ClpXP degradation assay using a more discriminatory set of time points, which revealed that the native CATssrA protein translated with the Shuf1 coding sequence is somewhat more susceptible to degradation than WT-CATssrA. This does not alter the main conclusions of this paper, that synonymous codon substitutions can affect protein folding in vivo to an extent that affects cell fitness. To the contrary: this new result provides direct confirmation of the impact of the synonymous codon changes on CAT protein folding. The manuscript text has been revised to improve the clarity of presentation of our results and conclusions.

